# ExpansionHunter: A sequence-graph based tool to analyze variation in short tandem repeat regions

**DOI:** 10.1101/572545

**Authors:** Egor Dolzhenko, Viraj Deshpande, Felix Schlesinger, Peter Krusche, Roman Petrovski, Sai Chen, Dorothea Emig-Agius, Andrew Gross, Giuseppe Narzisi, Brett Bowman, Konrad Scheffler, Joke J.F.A. van Vugt, Courtney French, Alba Sanchis-Juan, Kristina Ibáñez, Arianna Tucci, Bryan Lajoie, Jan H. Veldink, Lucy Raymond, Ryan J. Taft, David R. Bentley, Michael A. Eberle

**Affiliations:** Illumina Inc., 5200 Illumina Way, San Diego, CA, USA; Illumina Cambridge Ltd., Chesterford Research Park, Little Chesterford, UK; New York Genome Center, 101 Avenue of the Americas, New York, NY, USA; Department of Neurology, Brain Center Rudolf Magnus, University Medical Center Utrecht, Utrecht, The Netherlands; Department of Medical Genetics, University of Cambridge, Cambridge, UK; Department of Haematology, University of Cambridge, Cambridge, CB2 0PT, UK; NIHR BioResource, Cambridge University Hospitals NHS Foundation Trust, Cambridge, CB2 0QQ, UK; Genomics England, Queen Mary University London, Dawson Hall, London, EC1M 6BQ

## Abstract

We describe a novel computational method for genotyping repeats using sequence graphs. This method addresses the long-standing need to accurately genotype medically important loci containing repeats adjacent to other variants or imperfect DNA repeats such as polyalanine repeats. Here we introduce a new version of our repeat genotyping software, ExpansionHunter, that uses this method to perform targeted genotyping of a broad class of such loci.

**Availability and implementation:** ExpansionHunter is implemented in C++ and is available under the Apache License Version 2.0. The source code, documentation, and Linux/macOS binaries are available at https://github.com/Illumina/ExpansionHunter/.

## Introduction

Short tandem repeats (STRs) are ubiquitous throughout the human genome. Although our understanding of STR biology is far from complete, emerging evidence suggests that STRs play an important role in basic cellular processes (Hannan 2018; Gymrek et al. 2016). In addition, STR expansions are a major cause of over 20 severe neurological disorders including amyotrophic lateral sclerosis, Friedreich ataxia (FRDA), and Huntington’s disease (HD).

ExpansionHunter was the first computational method for genotyping STRs from short-read sequencing data capable of consistently genotyping repeats longer than the read length and, hence, detecting pathogenic repeat expansions (Dolzhenko et al. 2017). Since the initial release of ExpansionHunter, several other methods have been developed and were shown to accurately identify long (greater than read length) repeat expansions (Dashnow et al. 2018; Tang et al. 2017; Tankard et al. 2018; Mousavi et al. 2019).

Current methods are not designed to handle complex loci that harbor multiple repeats. Important examples of such loci include the CAG repeat in the *HTT* gene that causes HD flanked by a CCG repeat, the GAA repeat in *FXN* that causes FRDA flanked by an adenine homopolymer, and the CAG repeat in *ATXN8* that causes Spinocerebellar ataxia type 8 (SCA8) flanked by an ACT repeat. An even more extreme example is the CAGG repeat in the *CNBP* gene whose expansions cause Myotonic Dystrophy type 2 (DM2). This repeat is adjacent to polymorphic CA and CAGA repeats (Liquori et al. 2001) making it particularly difficult to accurately align reads to this locus. Another type of complex repeat is the polyalanine repeat which has been associated with at least nine disorders to date (Shoubridge and Gecz 2012). Polyalanine repeats consist of repetitions of α-amino acid codons GCA, GCC, GCG, or GCT (i.e. GCN).

Clusters of variants can affect alignment and genotyping accuracy (Lincoln et al. 2019). Variants adjacent to low complexity polymorphic sequences can be additionally problematic because methods for variant discovery can output clusters of inconsistently represented or spurious variant calls in such genomic regions. This, in part, is due to the elevated error rates of such regions in sequencing data (Benjamini and Speed 2012; Dolzhenko et al. 2017). One example is a single-nucleotide variant (SNV) adjacent to an adenine homopolymer in *MSH2* that causes Lynch syndrome I (Froggatt et al. 1999).

Here we present a new version (v3.0.0) of ExpansionHunter that was re-implemented to handle complex loci such as those described above. The implementation uses sequence graphs (Garrison et al. 2018; Paten et al. 2017; Dilthey et al. 2015) as a general and flexible model of each target locus.

## Implementation

ExpansionHunter works on a predefined variant catalog containing genomic locations and the structure of a series of targeted loci. For each locus, the program extracts relevant reads (Dolzhenko et al. 2017) from a binary alignment/map (BAM) file (Li et al. 2009) and realigns them using a graph-based model representing the locus structure. The realigned reads are then used to genotype each variant at the locus (Figure 1).

**Figure 1:**
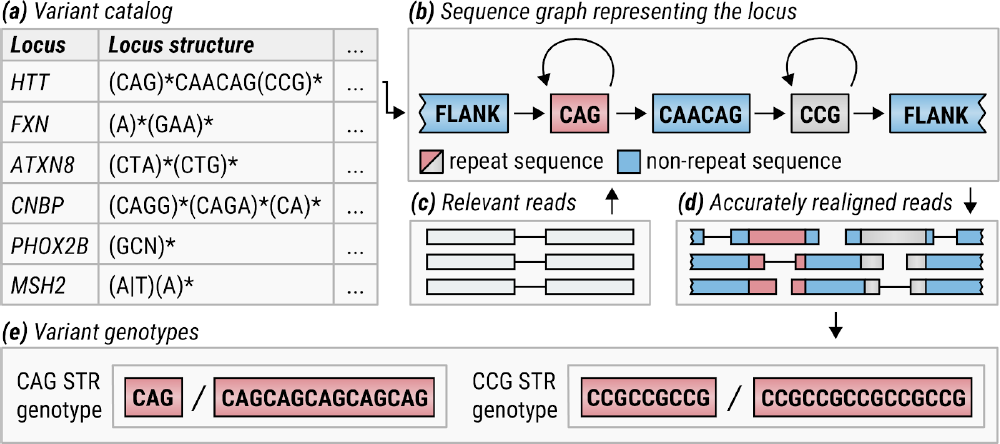
Overview of ExpansionHunter. (a) A locus definition is read from the variant catalog file. (b) Sequence graph is constructed according to its specification in the variant catalog. (c) Relevant reads are extracted from the input BAM file. (d) Reads are aligned to the graph. (e) Alignments are pieced together to genotype each variant.

The locus structure is specified using a restricted subset of the regular expression syntax. For example, the *HTT* repeat region linked to HD can be defined by expression (CAG)*CAACAG(CCG)* that signifies that it harbors variable numbers of the CAG and CCG repeats separated by a CAACAG interruption (see supplementary materials); the *FXN* repeat region linked to the FRDA corresponds to expression (A)*(GAA)*; the ATXN8 repeat region linked to SCA8 corresponds to (CTA)*(CTG)*; the *CNBP* repeat region linked to DM2 consists of three adjacent repeats is defined by (CAGG)*(CAGA)*(CA)*; the *MSH2* SNV adjacent to an adenine homopolymer that causes Lynch syndrome I corresponds to (A|T)(A)*.

Additionally, the regular expressions are allowed to contain multi-allelic or “degenerate” base symbols that can be specified using the International Union of Pure and Applied Chemistry (IUPAC) notation (“Nomenclature for Incompletely Specified Bases in Nucleic Acid Sequences. Recommendations 1984. Nomenclature Committee of the International Union of Biochemistry (NC-IUB)” 1986). Degenerate bases make it possible to represent certain classes of imperfect DNA repeats where, for example, different bases may occur at the same position. Using this notation, polyalanine repeats can be encoded by the expression (GCN)* and polyglutamine repeats can be encoded by the expression (CAR)*.

ExpansionHunter translates each regular expression into a sequence graph. Informally, a sequence graph consists of nodes that correspond to sequences and directed edges that define how these sequences can be connected together to assemble different alleles. We implemented the basic sequence graph functionality used by ExpansionHunter in the GraphTools C++ library (supplementary materials). One of the key features of the library is its support for single node loops in contrast to the traditional approaches that use fully acyclic graphs (Lee, Grasso, and Sharlow 2002). Single-node loops are the key to representing STRs and other sequences that can appear in any number of copies.

Genotyping is performed by analyzing the alignment paths associated with the presence or absence of each constituent allele. The repeats are genotyped as before (Dolzhenko et al. 2017) and SNVs/indels are genotyped using a straightforward Poisson-based model (supplementary materials).

## Results and discussion

To demonstrate the performance of ExpansionHunter we analyzed multiple complex STR regions. First, we analyzed a simulated dataset containing a wide range of CAG and CCG repeat sizes at the *HTT* locus. As expected, the accuracy of ExpansionHunter was substantially higher when the reads were aligned to a sequence graph that included both repeats compared to when the repeats were analyzed independently (Supplementary Figure S2). ExpansionHunter also produced more accurate genotypes compared to other tools that were not designed to handle loci harboring multiple nearby STRs, GangSTR and TREDPARSE (Supplementary Figure S2). A recent study used ExpansionHunter to investigate mutations in the short sequence interrupting two repeats in the *HTT* locus across 1,600 samples (Wright et al. 2019) demonstrating usefulness of the program for analysis of complex loci in real data. ExpansionHunter also correctly detected the pathogenic SNV adjacent to an adenine homopolymer in the *MSH2* gene in three WGS replicates of a sample obtained from SeraCare Life Sciences (Supplementary Materials).

To demonstrate the utility of ExpansionHunter across both short and long repeats, we compared calls from ExpansionHunter, GangSTR, and TREDPARSE on sequence data from samples with experimentally-confirmed repeat expansions (Supplementary Materials and Figure S3). ExpansionHunter had better accuracy (precision = 0.91, recall = 0.99) in detecting the expanded repeats in this dataset compared to GangSTR (precision = 0.88, recall = 0.83) and TREDPARSE (precision = 0.84, recall = 0.46).

Finally, we used ExpansionHunter to genotype degenerate DNA repeats by analyzing a polyalanine repeat in *PHOX2B* gene in 150 healthy controls and one sample harboring a known pathogenic expansion. *PHOX2B* contains a polyalanine repeat of 20 codons that can expand to cause congenital central hypoventilation syndrome. Consistent with what is known about this repeat (Amiel et al. 2003), all but a few controls were genotyped 20/20. ExpansionHunter accurately genotyped the sole sample with the expansion as 20/27; the correctness of this genotype was confirmed by Sanger sequencing.

In summary, we have developed a novel method that addresses the need for more accurate genotyping of complex loci. This method can genotype polyalanine repeats and resolve difficult regions containing repeats in close proximity to small variants and other repeats. A catalog of difficult regions is supplied with the software and can be extended by the user. We expect that the flexibility of the sequence graph framework now adopted in ExpansionHunter will enable a variety of novel variant calling applications.

