## Supplementary Methods for "ExpansionHunter: A sequence-graph based tool to analyze variation in short tandem repeat regions"

### Supplementary Materials

#### GraphTools library

We implemented the basic sequence graph functionality used by ExpansionHunter in the GraphTools library. This library implements core graph abstractions (graphs themselves, graph paths, and graph alignments), operations on them, and algorithms for aligning linear sequences to graphs.

In our implementation, a sequence graph consists of nodes and directed edges. The graphs are allowed to contain self-loops (an edge connecting a node to itself) but no other cycles. The nodes contain sequences made up of core base symbols and IUPAC degenerate base codes.

A graph path is defined by a sequence of nodes that the path goes through together with the start position of the path on the first node and the end position on the last node. The positions are specified using the zero-based half-open coordinate system. The library defines multiple operations on paths including path extension and shrinkage, overlap checks, and path merging.

Graph alignments encode how linear query sequences (usually sequenced reads) are aligned to the graphs. A graph alignment consists of a graph path and a sequence of linear alignments defining the alignment of the query sequence to each node overlapped by the path. Using the corresponding operations on paths, graph alignments can be shrunk or merged with other graph alignments. Path shrinking provides a mechanism for removing low confidence ends of the alignments while alignment merging is used by graph alignment algorithms to stitch together the full alignment of the query sequence from alignments of subsequences.

The alignment algorithm implemented in GraphTools operates by finding a kmer match between the query sequence and the graph, extracting a subgraph around the path corresponding to the kmer match (unrolling any cycles in the process). Then it performs a Smith–Waterman (Smith and Waterman 1981) alignment against the resulting acyclic graph. The algorithm supports affine gap penalties and is written using constant-length loops to enable compilers to generate SIMD code.

#### Application architecture

ExpansionHunter is designed as a general tool for targeted variant genotyping (Figure S1, left panel). During each run, the program attempts to genotype a set of variants described in the variant catalog file. The variants located in close proximity of each other are grouped into a single locus. The locus structure is specified using a restricted subset of the regular expression

(RE) syntax. REs contain sequences over the alphabet consisting of core base symbols and IUPAC degenerate base codes and must contain one or more of the expressions (<sequence>)?, (<sequence a>|<sequence b>), (<sequence>)*, (<sequence>)+ possibly separated by sequence interruptions. These expressions correspond to insertions/deletions, substitutions, sequence repeating 0 or more times, and sequence repeating at least once respectively. Figure S4 illustrates how a regular expression is transformed into a graph. Additionally, the description of each locus contains a set of reference regions for that locus and reference coordinates of each constituent variant.

The bulk of the work is orchestrated by objects of `LocusAnalyzer` class that synthesizes a sequence graph representing the locus from the corresponding RE during initialization. After initialization, a locus analyzer processes the relevant reads by aligning them to the graph and then passing the resulting alignments to `VariantAnalyzer` that is defined for each variant contained in the locus. A `VariantAnalyzer` extracts information relevant for genotyping the associated variant and passes it to the `VariantGenotyper` that performs the actual genotyping. The results output by each genotyper are then used to create the output VCF file.

For example, the `LocusAnalyzer` responsible for processing the locus containing a pathogenic variant associated with Lynch I syndrome utilizes SNV analyzer and STR analyzer (Figure S1, right panel).

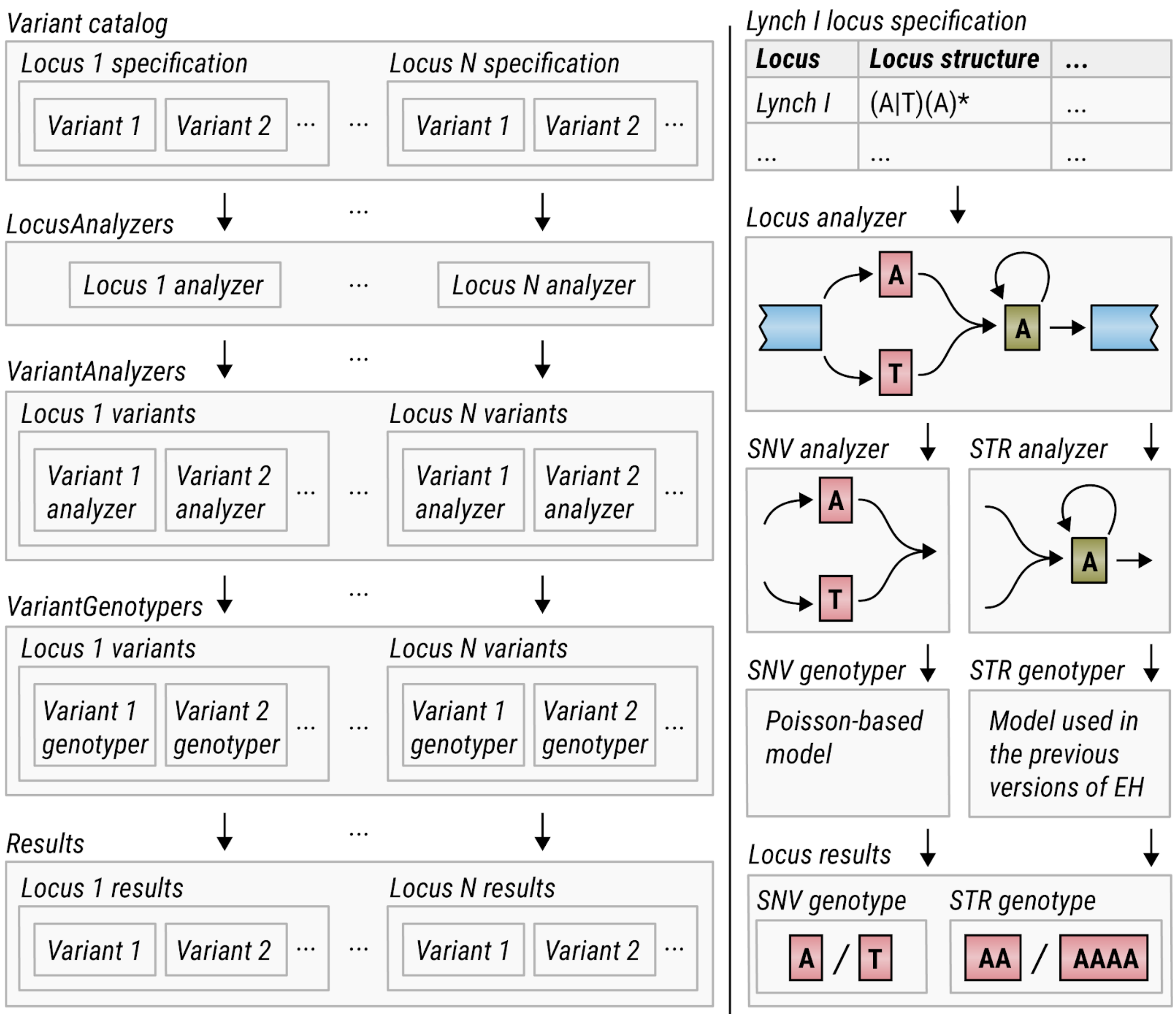


**Figure S1:** (left panel) High-level overview of the targeted genotyping framework implemented by ExpansionHunter; (right panel) application of the framework to the locus containing a pathogenic variant associated with Lynch I syndrome.

### Indel genotyper

Some STRs may have a short insertion, deletion, or substitution nearby. Such variants  are modeled as additional sub-graphs in the flanking sequences of the STR. The number of reads mapped to each allele (corresponding to a path in the sub-graph) is modeled with a Poisson distribution whose rate parameter is estimated from the mean read depth and read length observed at the locus. Genotype likelihoods are calculated under a Bayesian framework.

#### Realignment of reads to the graph

After a sequence graph representing the target locus is synthesized from the corresponding entry in the catalog file the program extracts reads from the input BAM file that are aligned to the corresponding reference region extended upstream and downstream by 1 Kb (by default). If any mates of the extracted reads align outside of this region they are also extracted and realigned. Finally, some repeats can be genotyped past the fragment length by including reads where both mates originate inside the repeat (in-repeat read pairs) (Dolzhenko et al. 2017). In-repeat read pairs are prone to mismapping and often both mates are misaligned far from the target locus in the input BAM file. If locations where such reads can mismap are known and specified in the catalog file, ExpansionHunter will also extract and realign reads from these regions. Note that in-repeat read pairs should only be included in the analysis when the user is confident that these reads are derived from the target repeat because including in-repeat read pairs for repeats that are common in the genome could lead to spurious false positives.

#### Definition of repeat regions

We specified all repeats in their reference orientation since this orientation is assumed by the program. For example, the *CNBP* repeat region is typically specified as (TG)*(TCTG)*(CCTG)* in the literature while we define at as (CAGG)*(CAGA)*(CA)* to be consistent with the reference genome.

Note also that there are other ways to define the *HTT* repeat region. For example, a more biologically sound definition would use 12 bp interrupting sequence between the pure CAG and CCG repeats instead of the 6pb interruption used here. This definition would increase the similarity between STR and the interrupting sequence and hence could result in loss of genotyping accuracy making this definition less robust from a computational perspective.

#### Analysis of CAG and CCG STRs in the *HTT* locus

We simulated short read samples for a wide range of CAG and CCG repeat sizes in the *HTT* locus using wgsim (Li, n.d.). We set the read length to 150, distance between mate ends to 350, standard deviation for mate end distance to 50, rate of mutations and base error rate to 0.0010, and fraction of indels to 0. The number of pairs was set to yield 40x coverage of the locus. The reads were aligned to GRCh37 reference with BWA-MEM (Li 2013).

To demonstrate the increased power provided by the sequence graph genotyping model, we analyzed the resulting dataset with ExpansionHunter in two ways. First, we specified the structure of the HTT locus using the expression (CAG)*CAACAG(CCG)* ensuring that reads are aligned to the sequence graph containing both repeats. Then we used ExpansionHunter to analyze each repeat independently. In this mode, the reads were aligned to the graph that only included the CAG STR and, separately, to the graph that only included the CCG STR. Finally we analyzed both repeats independently with GangSTR v2.3 and TREDPARSE (commit c856d5c) (Figure S2).

ExpansionHunter accurately estimated genotypes when reads were aligned to a graph containing both repeats compared to separately genotyping each repeat with either of the three tools. This showcases the benefits of accurately modeling the structure of a region with sequence graphs.

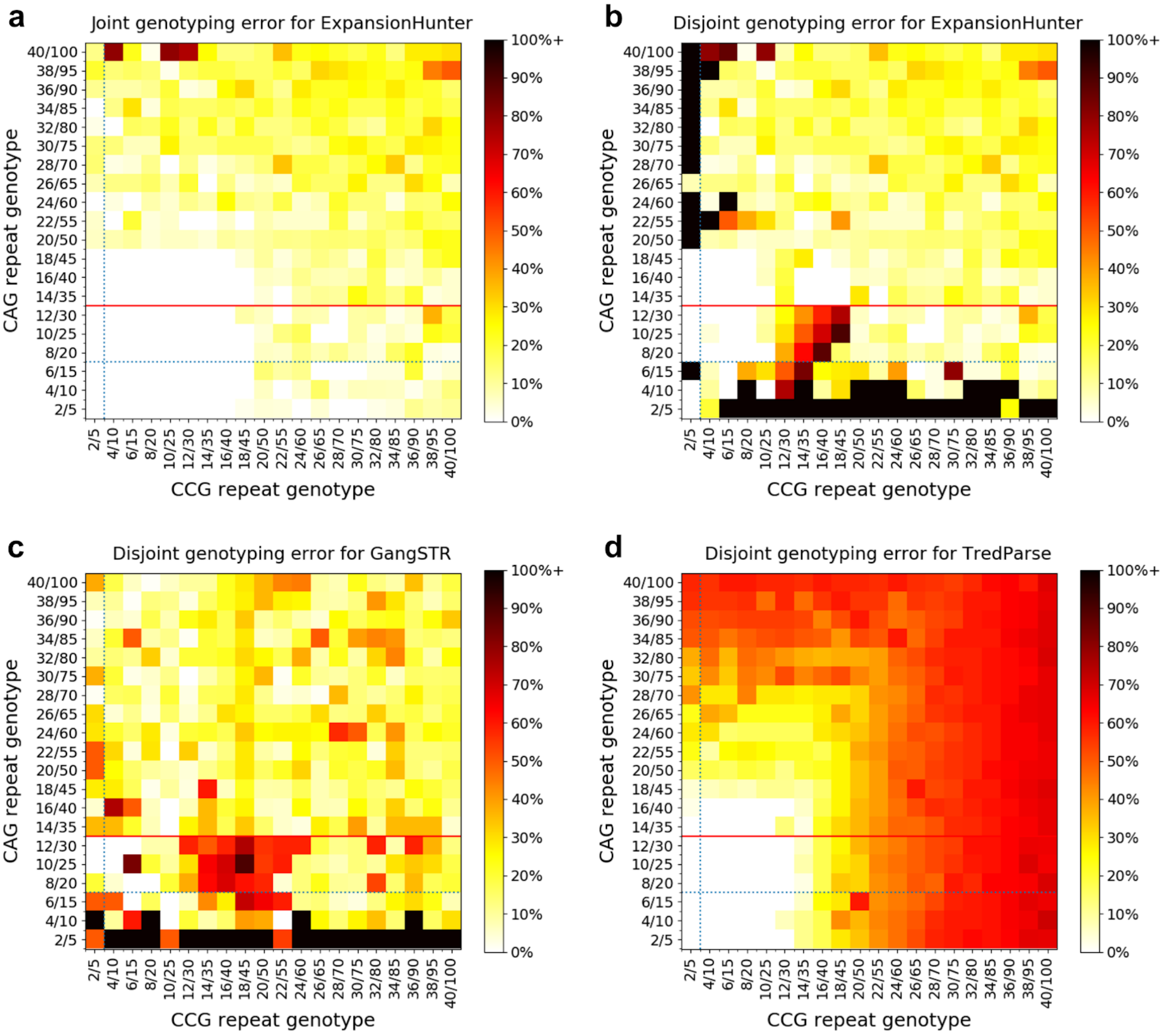


**Figure S2:** Accuracy of genotyping CAG and CCG STRs in the *HTT* locus from simulated data. (a) Performance of ExpansionHunter when reads are aligned to sequence graph containing both repeats. Performance of (b) ExpansionHunter, (c) GangSTR and (d) TREDPARSE when the repeats are analyzed independently. For each simulated sample, we measured the maximum percentage deviation of the predicted STR lengths from the expected STR lengths.

Dotted blue lines correspond to the genotype in the reference genome and solid red lines represent the threshold for pathogenic expansions.

#### Analysis of samples with experimentally-confirmed expansions

We analyzed 120 samples from Coriell (Dolzhenko et al. 2017) harboring repeat expansions associated with dentatorubral-pallidoluysian atrophy (ATN1 gene), fragile X Syndrome (FMR1 gene), Friedreich's ataxia (FXN gene), Huntington's disease (HTT gene), myotonic dystrophy type 1 (DMPK gene), spinocerebellar ataxia type 1 (ATXN1 gene), spinocerebellar ataxia type 3 (ATXN3 gene), and spinal and bulbar muscular atrophy (AR gene).

The results from ExpansionHunter, GangSTR, and TREDPARSE analyses are shown in Figure S3. GangSTR was run only on STRs for which predefined off-target loci were provided by the authors so we restricted our recall and precision calculations to these STRs for comparison. We classified each genotype according to whether the corresponding tool estimates that the size of the longest allele is above or below the threshold for normal repeats. This classification yielded the following accuracy estimates: precision of 0.91 and recall of 0.99 for ExpansionHunter; precision of 0.88 and recall of 0.83 for GangSTR; precision of 0.84 and recall of 0.46 for TREDPARSE. Note also that we ran GangSTR for *FMR1* repeat in targeted mode and in the genome-wide mode for the other repeats. This was done because the results were more accurate for *FMR1* and the other repeats in these respective modes. For comparison, the precision and recall were 0.90 and 0.52 using just the genome-wide mode and 0.22 and 0.86 using just the targeted mode.

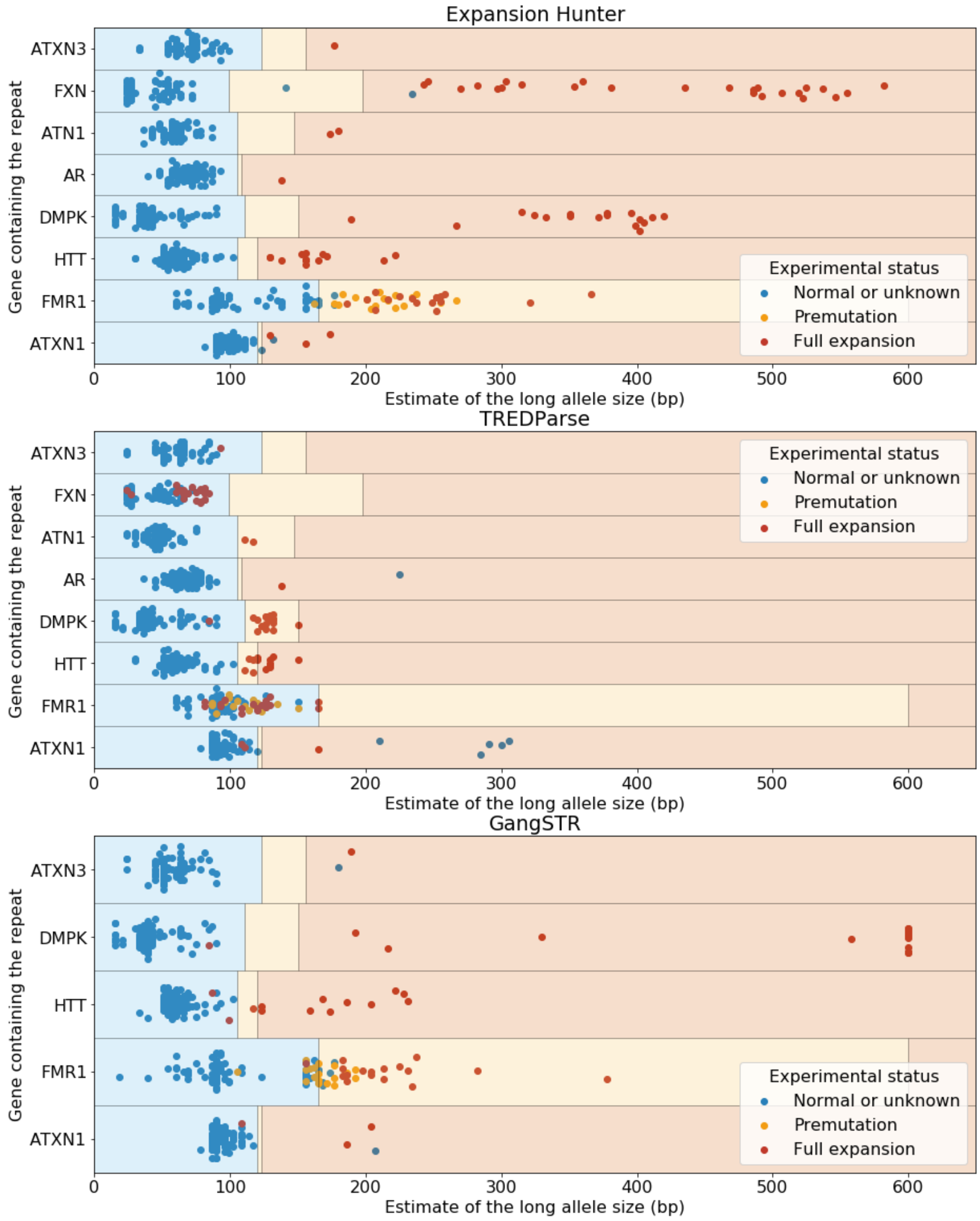


**Figure S3:** Analysis of Coriell samples harboring known repeat expansions. The blue, orange, and red rectangles define the expected size ranges for normal, premutation, and full expansion respectively for the corresponding repeat. Each dot corresponds to the size of the longest allele and its color is set according to the experimentally-determined status. GangSTR was run only on STRs for which predefined off-target loci were provided. GangSTR values were calculated using their “genome-wide” mode for all of the genes except *FMR1* which was analyzed using “targeted” mode which performed much better for this repeat. The repeat sizes were capped at 600bp.

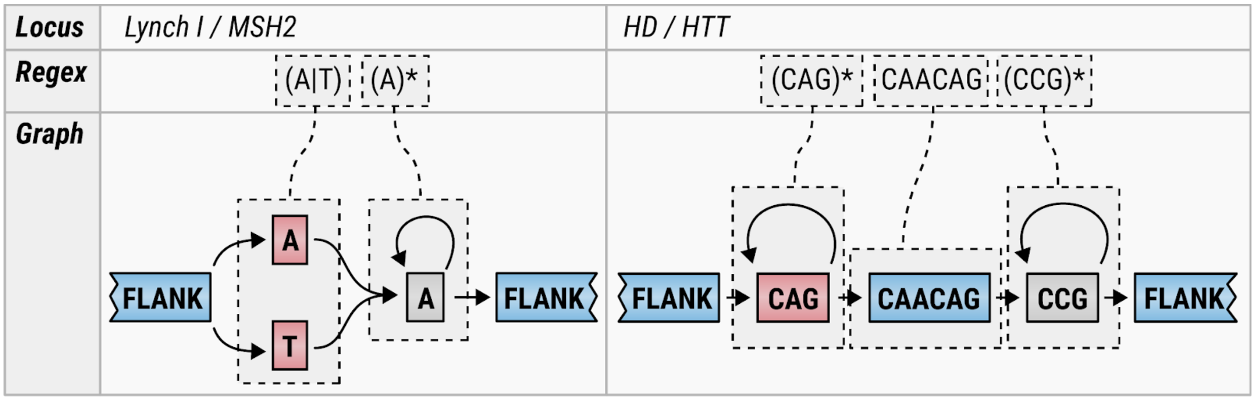


**Figure S4:** Examples illustrating the correspondence between regular expression and sequence graphs.

#### Detection and annotation of regions containing multiple polymorphic STRs

To obtain a rough lower bound on the number of complex STR loci in the genome we performed a simultaneous analysis of alignments from 25 unrelated samples. We searched the regions within 5kbp of ~19,000 gene bodies where read alignments from multiple samples contained insertions and deletions consistent with presence of a polymorphic STR region. As illustrated in Figure S5, these insertions and deletions consist of repetitions of some repeat unit.  So far, our search revealed 56,858 polymorphic STR loci. Of these, 2,879 loci (5%) contained two or more adjacent STRs. Given the difficulty of aligning reads to these regions we expect that our analysis significantly under represents the fraction of complex loci. More development remains to be done to perform a more comprehensive search and to produce a high-quality STR catalog from these regions.

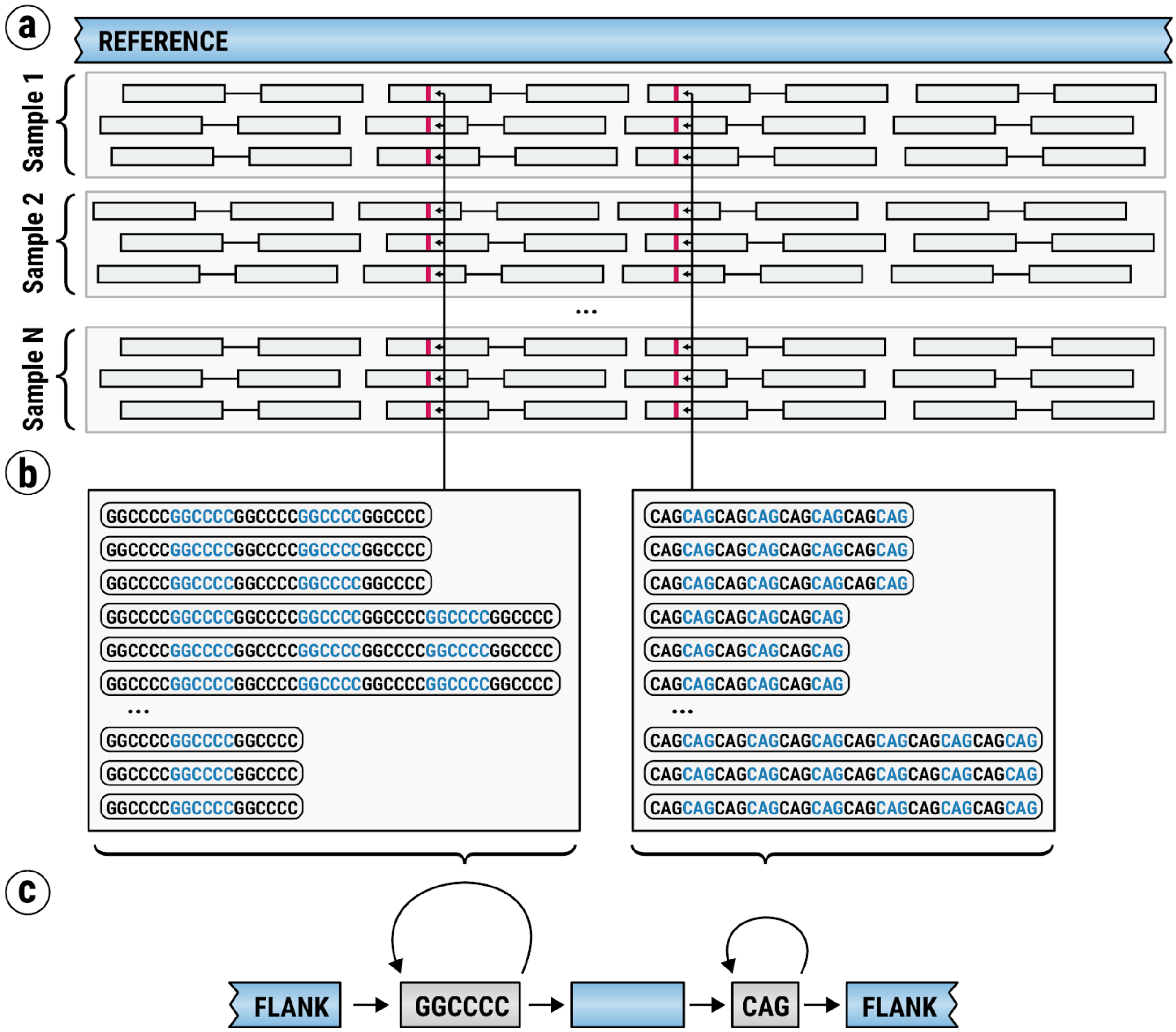


**Figure S5:** An overview of an approach to detect and annotate regions containing multiple polymorphic STRs. (a) Alignment of reads from N samples to a short region in the reference genome; indels in the alignments marked by vertical red bars. (b) Indel sequences from two nearby sites; indels at the first site consist of repetitions of GGCCCC hexamers and indels at the second site consist of repetitions of CAG trimers. (c) Sequence graph capturing the population-scale structure of the locus.

#### Using sequence graphs to model local versus genome-wide variation

The method described here uses sequence graphs to model local variation. A sequence graph synthesized from the input locus definition is used as a replacement for a small piece of reference enabling accurate genotyping of the constituent variants. This approach works well for STRs because the vast majority of relevant reads either map to the target locus in the input BAM file or have a mate that maps nearby making these reads easy to recover. Even the read pairs originating from very large repeat expansions where both mates are mismapped far from the target locus can be recovered with relatively straightforward methods (Dolzhenko et al. 2017).

At the same time, the approach presented in this work is not appropriate for analyzing larger scale variation. The vast majority of reads originating in, say, larger insertions not present in the linear reference will be misaligned or unaligned. For larger scale variants, a truly end-to-end solution will require alignment against a comprehensive pangenome reference such as is implemented in vg (Garrison et al. 2018).

#### Using general variant detection software for STR calling

To check the ability of general-purpose variant detection software to call STRs, we simulated a dataset containing a range of CAG repeat sizes surrounded by sequence flanking the pathogenic CAG repeat in the *HTT* locus. (The 2x150 reads were simulated with wgsim to 40x coverage depth of the locus and then aligned with bwa mem (Li 2013) to GRCh37 reference.) The resulting dataset was analyzed with FreeBayes (Garrison and Marth 2012) and ExpansionHunter (Figure S6). Both methods were able to accurately detect/genotype STRs significantly shorter than the read length. As expected, ExpansionHunter was also able to genotype longer STRs.

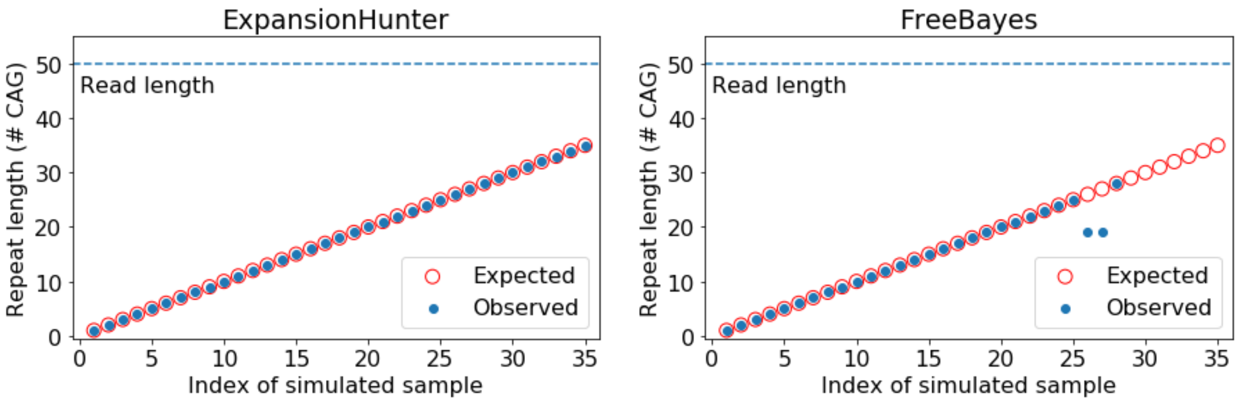


**Figure S6:** Analysis of simulated samples containing a range of sizes of the pathogenic CAG repeat in the *HTT* locus with ExpansionHunter and FreeBayes.

### Datasets

The PCR-free WGS sequencing data for 150 unrelated controls used for the analysis of the *PHOX2B* polyalanine repeat is the Polaris Diversity Cohort. It consists of samples selected from the International Genome Sample Resource (1000 Genomes Project Consortium et al. 2015) (http://www.internationalgenome.org/). The WGS data can be obtained from European Nucleotide Archive (ENA; https://www.ebi.ac.uk/ena/home; PRJEB20654) and from NCBI Sequence Read Archive (SRA; https://www.ncbi.nlm.nih.gov/sra; bioproject:387148). The description of the samples is available online (https://github.com/Illumina/Polaris/wiki/HiSeqX-Diversity-Cohort). The PCR-free WGS sequence data for the 120 samples with known pathogenic repeats is available in the European Genome-phenome Archive (EGA; https://www.ebi.ac.uk/ega/home; EGAD00001003562).

The sample with the 20/27 expansion in *PHOX2B* was obtained from the Genetics Laboratories Molecular Genetics, Addenbrooke's Treatment Centre, Cambridge University.

The SeraCare Life Sciences sample, Seraseq Inherited Cancer DNA Mix v1, contains variants that are known to be both pathogenic and difficult-to-call. The variants have been synthetically added to the well-characterized cell line GM24385, all with expected variant frequencies of 50%. One of these engineered mutations corresponds to an SNV in the *MSH2* gene, which is directly adjacent to a long homopolymer A region.

Three replicates of the SeraCare sample were prepared with the Illumina TruSeq PCR Free kit. The replicates were run on a single lane of the NovaSeq6000 using the XP workflow and sequenced at 2x151 read length. The replicates were analyzed using the 'Sentieon DNASeq FASTQ to VCF' and 'Whole Genome Resequencing v8.0.0' Basespace apps (https://basespace.illumina.com). Sentieon can be regarded as a proxy for variant calling performance of the Broad's BWA-GATK software suite as Sentieon implements the same algorithms. Neither of the software solutions was able to correctly identify the *MSH2* SNV. Fastq files and analysis results are provided through Basespace: https://basespace.illumina.com/s/HAQNxJyEtJLP


1000 Genomes Project Consortium, Adam Auton, Lisa D. Brooks, Richard M. Durbin, Erik P. Garrison, Hyun Min Kang, Jan O. Korbel, et al. 2015. "A Global Reference for Human Genetic Variation." *Nature* 526 (7571): 68–74.

Dolzhenko, Egor, Joke J. F. A. van Vugt, Richard J. Shaw, Mitchell A. Bekritsky, Marka van Blitterswijk, Giuseppe Narzisi, Subramanian S. Ajay, et al. 2017. "Detection of Long Repeat Expansions from PCR-Free Whole-Genome Sequence Data." *Genome Research* 27 (11): 1895–1903.

Garrison, Erik, and Gabor Marth. 2012. "Haplotype-Based Variant Detection from Short-Read Sequencing." *arXiv [q-bio.GN]*. arXiv. http://arxiv.org/abs/1207.3907.

Garrison, Erik, Jouni Sirén, Adam M. Novak, Glenn Hickey, Jordan M. Eizenga, Eric T. Dawson, William Jones, et al. 2018. "Variation Graph Toolkit Improves Read Mapping by

Representing Genetic Variation in the Reference.” *Nature Biotechnology* 36 (9): 875–79.
Li, Heng. 2013. “Aligning Sequence Reads, Clone Sequences and Assembly Contigs with BWA-MEM.” *arXiv [q-bio.GN]*. arXiv. http://arxiv.org/abs/1303.3997.
———. n.d. “Tool for Simulating Sequence Reads from a Reference Genome.” https://github.com/lh3/wgsim.
Smith, T. F., and M. S. Waterman. 1981. “Identification of Common Molecular Subsequences.” *Journal of Molecular Biology* 147 (1): 195–97.
